# Menstrual cycle associated alteration of vastus lateralis motor unit function

**DOI:** 10.1101/2023.03.27.534396

**Authors:** Jessica Piasecki, Yuxiao Guo, Eleanor J. Jones, Bethan E. Phillips, Daniel W. Stashuk, Philip J. Atherton, Mathew Piasecki

## Abstract

Estrogen and progesterone are the primary female sex hormones and have net excitatory and inhibitory effects, respectively, on neuronal function. Fluctuating concentrations across the menstrual cycle has led to several lines of research in relation to neuromuscular function, yet evidence from animal and cell culture models have yet to be demonstrated in human motor units (MU) coupled with quantification of circulating hormones.

Intramuscular electromyography (iEMG) was applied to record MU potentials (MUP) and corresponding MUP trains (MUPT) from the vastus lateralis of eumenorrheic females during the early follicular, ovulation and mid luteal phases of the menstrual cycle, alongside assessments of neuromuscular performance. Multi-level regression models were applied to explore effects of time and of contraction level. Statistical significance was accepted as p<0.05.

Knee extensor maximum voluntary contraction (MVC), jump power, force steadiness, and balance did not differ across the menstrual phases (all p>0.4). Firing rate of low threshold MU (10% MVC) was reduced during phases of high progesterone (β=-0.82Hz, p<0.001), with no difference in MUPs analysed from 25% MVC contractions. MUPs were more complex during ovulation and mid luteal phase (p<0.03), with no change in neuromuscular junction transmission instability (p>0.3).

Assessments of neuromuscular performance did not differ across the menstrual cycle. The suppression of low threshold MU firing rate during periods of increased progesterone may suggest a potential inhibitory effect and an alteration of recruitment strategy, however this had no discernible effect on performance. These findings highlight contraction level dependent modulation of VL MU function over the eumenorrheic cycle.

## Introduction

Estrogen and progesterone are the primary female sex hormones and have the ability to cross the blood-brain barrier and potentially influence the functionality of the central nervous system (CNS) (Stoffel-Wagner, 2001). The fluctuating concentrations of these hormones characterises the menstrual cycle (changing over a period of 22-35 days (Fehring, Schneider and Raviele, 2006)) which is a natural process for most biological females and serves several physiological maintenance roles beyond reproduction. Although a highly heterogeneous process (Bull *et al*., 2019; Bruinvels *et al*., 2021a), circulating estrogen typically rises to a peak around day 10-14 followed by a slow decrease over the following 5 days (Messinis, Messini and Dafopoulos, 2014). The latter 14-day luteal phase is characterised by a gradual increase in progesterone, peaking around day 22 and returning to base levels at day 28 (Messinis, Messini and Dafopoulos, 2014). It is unsurprising the regularly fluctuating concentrations of these hormones with genomic and nongenomic effects has attracted notable research interest in terms of general health (Eshima *et al*., 2019; Schmalenberger *et al*., 2019; Shayani *et al*., 2020; Bruinvels *et al*., 2022), neuromuscular performance (Tenan *et al*., 2013; Tenan, Hackney and Griffin, 2016; Ansdell *et al*., 2019; Weidauer *et al*., 2020), and injury risk (Wojtys *et al*., 1998; Shultz *et al*., 2004; Ekenros *et al*., 2017; Martin *et al*., 2021).

Estrogen receptors (ERs) are abundant in the CNS and estrogen elicits primarily net excitatory effects via several mechanisms. The ER-α binding of estradiol (E2), the most active type of estrogen, on GABAergic neurons initiates the suppression of GABA (the primary inhibitory neurotransmitter) via destabilisation of GABA receptors minimising inhibitory action in cultured neurons and hippocampal sections (Mukherjee *et al*., 2017). E2 supports the potentiation of the effects of excitatory glutamatergic neurons (Smith and Woolley, 2004), and promotes glutamate receptor trafficking (Potier *et al*., 2016). E2 has also been shown to affect the signalling of the neurotransmitter serotonin throughout the CNS which may have downstream effects on motoneuron firing rates (Michopoulos, Berga and Wilson, 2011; Bethea and Reddy, 2012; Barth, Villringer and Sacher, 2015). Opposingly, progesterone has a net inhibitory effect, shown to cause a decrease in discharge rate of Purkinje neurons during locomotion in animal models (Smith, Woodward and Chapin, 1989) and increased inhibition of rat pyramidal neurons (Hsu and Smith, 2003). Progesterone also reduces the availability of estrogen specific receptors on neuronal cells and enhances the inhibitory responses of limbic neurons (Smith and Woolley, 2004), supresses the excitatory response of glutamate (Smith *et al*., 1987) and increases GABA release (Smith, 1994; Bitran, Shiekh and McLeod, 1995; Melcangi *et al*., 2014). Although direct excitatory and inhibitory effects of these hormones have been well described, it is important to note a net neuronal effect in complex physiological systems (e.g. human performance) is a result of factors beyond estrogen and progesterone alone.

In addition to influencing neural excitability, estrogen may also influence neuromuscular performance directly at the muscle fibre, potentially enhancing myosin binding (Lowe, Baltgalvis and Greising, 2010). In smooth muscle estrogen can acutely inhibit L-type Ca^2+^ channels and limit contractility (Wray and Noble, 2008), which are also located at pre and post-synaptic regions of the neuromuscular junction (NMJ) and may exert similar inhibitory effects there via interference of acetylcholine release and receptor binding (Giovannini *et al*., 2002). Motor unit (MU) firing rate is a key determinant of force generating capacity and may be susceptible to the inhibitory/excitatory effects of sex hormones. Of the only study to investigate this, alterations of vastus medialis MU initial discharge rate were reported across 5 stages of the menstrual cycle, which may have corresponded with fluctuating estrogen and progesterone (Tenan *et al*., 2013). Here the authors postulated progesterone may have the more dominant effect on MU firing, although this was not quantified (Tenan *et al*., 2013). Collectively, these studies provide strong indications of the acute excitatory and inhibitory effects of sex hormones.

Although physiologically plausible, the lack of *in-* or *ex-vivo* data at central and peripheral sites of the motor system is further complicated by the equivocal findings of the menstrual cycle on neuromuscular performance, with some reporting greater strength at the menstrual phase (Davies, Elford and Jamieson, 1991), the follicular phase (Phillips *et al*., 1996), at the mid-point of the cycle (Sarwar, Niclos and Rutherford, 1996; Tenan, Hackney and Griffin, 2016), and some reports of increases at the luteal phase (Birch and Reilly, 2002). Several others report no change across the cycle (Dibrezzo, Fort and Brown, 1988; Greeves *et al*., 1997; Janse De Jonge *et al*., 2001; Elliott *et al*., 2003; Kubo *et al*., 2009), yet 36 - 51% of athletes identify their menstrual symptoms to adversely affect their performance (Bruinvels *et al*., 2016; Martin *et al*., 2018). Importantly, recent systematic review and meta-analyses have highlighted a lack of consistent high quality research, alongside a largely trivial difference in strength across a eumenorrheic cycle (Blagrove, Bruinvels and Pedlar, 2020; McNulty *et al*., 2020).

There are currently no data quantifying MU adaptation combined with menstrual tracking and hormone quantification across the human menstrual cycle. Therefore, the purpose of the present study was to quantify the vastus lateralis (VL) MU function using intramuscular electromyography (iEMG) at different contraction intensities over three key stages: early follicular, ovulation, and mid luteal phases of a single menstrual cycle in healthy eumenorrheic young females.

## Methods

### Ethical Approval

The research study was approved by local ethics committees at the University of Nottingham (302-1903). The study conformed with all standards set by Declaration of Helsinki, except for registration in a database. A total of 13 recreationally active females volunteered to take part in this research study. Three participants were unable to attend the second visit within the required ovulation window, and one participant was excluded based on low and non-changing progesterone levels. Full data are presented for nine females with a mean (SD) age of 24.2 (3.2) years and a BMI of 22.8 (2.8). Exclusion criteria included a diagnosis of metabolic disease, lower limb musculoskeletal abnormalities, acute cerebrovascular or cardiovascular disease, active malignancy, uncontrolled hypertension or those on medications that are potentially neuroactive or modulate vascular tone. All participants were classified as eumenorrheic having reported a regular cycle between 21-35 days (Fehring, Schneider and Raviele, 2006) for the previous 12 months, with a minimum of 9 cycles, and had not taken any oral contraceptive pill within 12 months. Participants arrived at the lab after an overnight fast and body mass and height were measured on the first visit only. The three testing sessions occurred at the same time of day for each stage of the cycle, and participants refrained from strenuous exercise and alcohol for 24 hours prior. To consider the influence of the menstrual cycle on neuromuscular performance participants were assessed at three time points across the cycle; 1) Early follicular; within 48 hours of the onset of the menstrual period. 2) at ovulation; determined by home based ovulation kits used to detect the rise in luteinizing hormone associated with ovulation. Upon a positive test, participants were tested in the lab within 48 hours. 3) The mid luteal phase; assessed 7 days following ovulation.

### Plasma hormones

Two EDTA tubes (20ml) of blood were drawn from the antecubital vein at each visit. All tubes were centrifuged at 3200*g at 4°C for 20 minutes. Plasma was then aliquoted into 1ml ependorph tubes and stored at -80 °C for future analysis. Stages of the menstrual cycle were confirmed via determination of the plasma concentration of 17β-estradiol and progesterone. Analysis of plasma samples was conducted using enzyme linked immunoassay kits (Invitrogen, Thermofisher Scientific, Camarillo, CA) following manufacturer’s instructions, measured at an absorbance of 450 nm. The minimal detection concentration of 17β-estradiol was 5 pg.mL and Progesterone was 4 pg.mL. Each sampled was added to the ELISA panel in duplicate, and a standard curve plotted with 6 standards for 17β-estradiol and 8 standards for progesterone. The concentration of hormones within the plasma samples was then determined from the mean absorbance of the duplicates, interpolating directly from the standard curve. The CV for 17β-estradiol was 3.9-6.1 % and for Progesterone 3.5-7.0 %. Participant hormonal samples all met previously recommended criteria (de Jonge, Thompson and Ahreum, 2019) when conducting experimental procedures across the menstrual cycle, with a peak in progesterone concentration at the mid-luteal phase and a rise in 17 β-estradiol from early follicular to ovulation. One participant had a progesterone concentration of 0.9 – 1.0 ng.ml at all time points and was not included in final analysis.

### Neuromuscular performance

Participants were seated in a custom-built chair with hips and knees flexed at 90°. The lower leg of the right limb was securely attached to a force dynamometer with non-compliant straps (purpose-built calibrated strain gauge, RS125 Components LTD, Corby, UK) above the medial malleolus. Participants were also strapped into the chair with a seat belt across the pelvis to minimise movement of the upper trunk throughout testing. All participants underwent a standardised warm up of submaximal isometric contractions. Once prepared and instructions fully understood, participants were then instructed to perform a maximum voluntary contraction (MVC), receiving visual feedback and verbal encouragement from the researchers. This was repeated a total of 3 times, with 60 seconds rest between each repetition, taking the highest values, in Newtons, to be the accepted MVC.

Prior to the force steadiness assessments, participants conducted a familiarisation, matching a target force on a screen, for around 12-15 seconds each time, at 10%, 25% and 40% MVC. Once familiarised participants were instructed to perform the experimental contractions, 4 sustained contractions at 10 and 25% MVC, and 2 at 40% MVC for 12-15 seconds each, with 30 seconds rest between each contraction. Force steadiness was quantified as the coefficient of variation of the force [CoV; (SD/mean) × 100]. To account for corrective actions when reaching the target line, the first two passes of the target (<1s) were excluded from the calculation. The mean CoV at each contraction level was calculated from the middle two contractions of 10 and 25%, and from both contractions at 40%.

An RS Foot scan (Gait and Motion Technology Ltd, Bury St Edmunds, UK) pressure sensor plate was used to assess single leg balance. The centre of pressure (COP), or postural sway, of the vertical plane was measured throughout assessment and is expressed as total distanced moved in millimetres (mm). This was assessed for 30 s with participants standing in the centre of the sensor plate, on the right leg only. To assess jump power, a G-walk sensor was placed at the base of the spine, secured with a Velcro waist belt (Gait and Motion Technology Ltd). Participants were asked to perform a counter movement jump, instructions were to jump as high as possible, with hands remaining on their waist with a trained assistant present and in reach of the participants in case of a fall or falter. Each participant repeated the jump sequence three times, with approximately 30 s rest between jumps, and the highest value was recorded (Piasecki *et al*., 2020).

### Intramuscular Electromyography (iEMG)

A 25 mm disposable concentric needle electrode (Ambu Neuroline, Ambu, UK) was used for all iEMG assessments. The needle was inserted into the muscle belly of the VL in the area of the motor point, established as previously described (Guo *et al*., 2022). iEMG signals were recorded using Spike 2 (Version 9.06) sampling at 50 KHz and bandpass filtered at 10 Hz to 10 kHz (1902 amplifier; Cambridge Electronic Design Ltd, Cambridge UK) and stored for future off-line analysis. A ground electrode was placed over the patella. Prior to sampling, participants were instructed to perform a series of voluntary low-level contractions with the needle in place to ensure adequate signal-to-noise ratio (e.g. visible spikes). Participants then performed sustained voluntary contractions as detailed above. After performing contractions at each contraction level the needle electrode was repositioned with combinations of turning the bevel 180 degrees and withdrawing by ∼5mm. This was repeated to perform 4 contractions recording from spatially distinct areas (Jones *et al*., 2021). Participants had 30 seconds rest between each contraction.

### iEMG Analysis

Decomposition-based quantitative electromyography (DQEMG) software was used to detect motor unit potentials (MUPs) and their corresponding motor unit potential trains (MUPTs). MUPTs composed of MUPs from more than one MU or with fewer than 30 MUPs were excluded from further analysis. Template MUPs of all MUPTs were visually checked, with markers adjusted where necessary to correspond to the onset, end, and positive and negative peaks. All MU data was analysed from the sustained phase of the contraction only, excluding rise and fall phases. iEMG data are reported for 10% and 25% MVC contractions only as reliable MUP identification was not possible for all participants at 40% MVC.

MU Firing rate (FR) was assessed as the rate of MUP occurrences within a MUPT, which is expressed as the number of occurrences per second (Hz). The MU FR variability is reported as the CoV for the interspike interval (ISI). Area of the MUP was measured as the total area within the MUP duration (from onset to end). MUP complexity is reported as the number of turns, measured as a change in waveform direction of at least 25 µV, indicating the level of temporal dispersion across individual muscle fibre contributions to a single MUP. A near fibre MUP (NFM) is defined as the low-pass-second derivative (Piasecki, Garnés-Camarena and Stashuk, 2021) of its corresponding MUP and is primarily contributed to by fibres within 350 µm of the recording surface of the electrode. This ensures that only potentials from fibres that are nearest to the needle electrode contribute to a NFM and that NFMs are composed of reduced contributions from more distant MU fibres and less interference from distant active fibres of other MUs. Any NFMs that did not have clear peaks were excluded from subsequent analysis. NFM jiggle is a measure of the variability of the shape of consecutive NFMs of an MUPT reflective of NMJ transmission instability, and are expressed as a percentage of the total NFM area (Piasecki, Garnés-Camarena and Stashuk, 2021).

### Statistical Analysis

All statistical analysis were performed using RStudio (V2022.07.1). Univariate ANOVA was performed to identify any differences between hormonal levels, MVC, jump power, balance performance and force steadiness. For MU characteristics, multi-level mixed-effect linear regression analysis was performed using the *lme4* package (V 1.1.23) (Bates *et al*., 2015). Models were generated to compare the three timepoints across the menstrual cycle at each contraction level, and then separately to compare differences with increasing contraction level (i.e from 10 to 25 % MVC) across stages. The linear mixed-effect modelling is appropriate for data of this nature as it incorporates the whole sample of extracted MUPTs, rather than just the mean values that were obtained from each participant, and provides coefficient estimates that indicate magnitude and direction of effects of interest. Results of these outputs are displayed as β coefficient estimates, 95% confidence intervals and p-values. Standardised betas and 95% confidence intervals are also shown for each model output. To explore the association between sex hormones and MU characteristics, separate linear models were generated for estrogen and progesterone, with participants means for each MU characteristic, and participant and time included as covariates. Statistical significance was assumed when p<0.05.

## Results

Estrogen concentrations differed across the 3 time points (p=0.002), with a mean of 244 pg.ml in the early follicular phase, rising to 533 pg.ml in the ovulatory phase (p=0.016). At 484 pg.ml in the mid-luteal phase, this was significantly higher than the early follicular (p=0.013) but not the ovulatory phase (p=0.744) (Figure 1A). Progesterone also differed across the 3 time points (p<0.001); with a mean of 2.66 ng.ml in the early follicular phase, rising to 3.22 ng.ml in the ovulatory phase (p=0.027), and rising again to 5.89 ng.ml in the mid-luteal phase (p=0.008), which was significantly higher than the early follicular phase (p<0.001) (Figure 1B).

**Figure 1.**
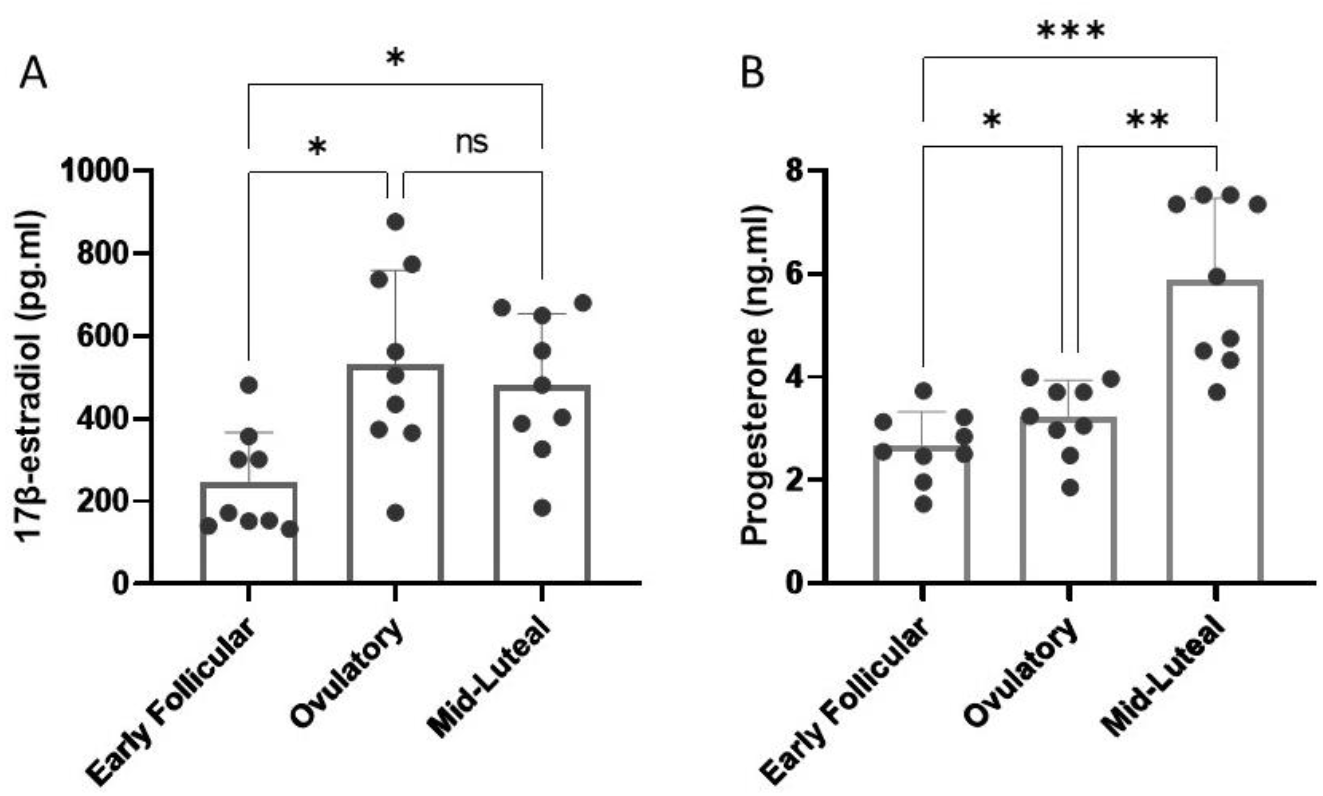
Circulating concentrations of A) estrogen (17β-estradiol) and B) progesterone across menstrual cycle phases. * p<0.05; ** p<0.01; ***p<0.001

There were no significant time associated differences in MVC (p=0.624), jump power (p=0.775), or distance travelled during right leg unilateral balance (p=0.713) (Figure 2A-C). Similarly, force steadiness did not differ across the cycle when assessed at 10% MVC (p=0.453), 25% (p=0.405), or 40% MVC (p=0.633) (Figure 2D-F).

**Figure 2.**
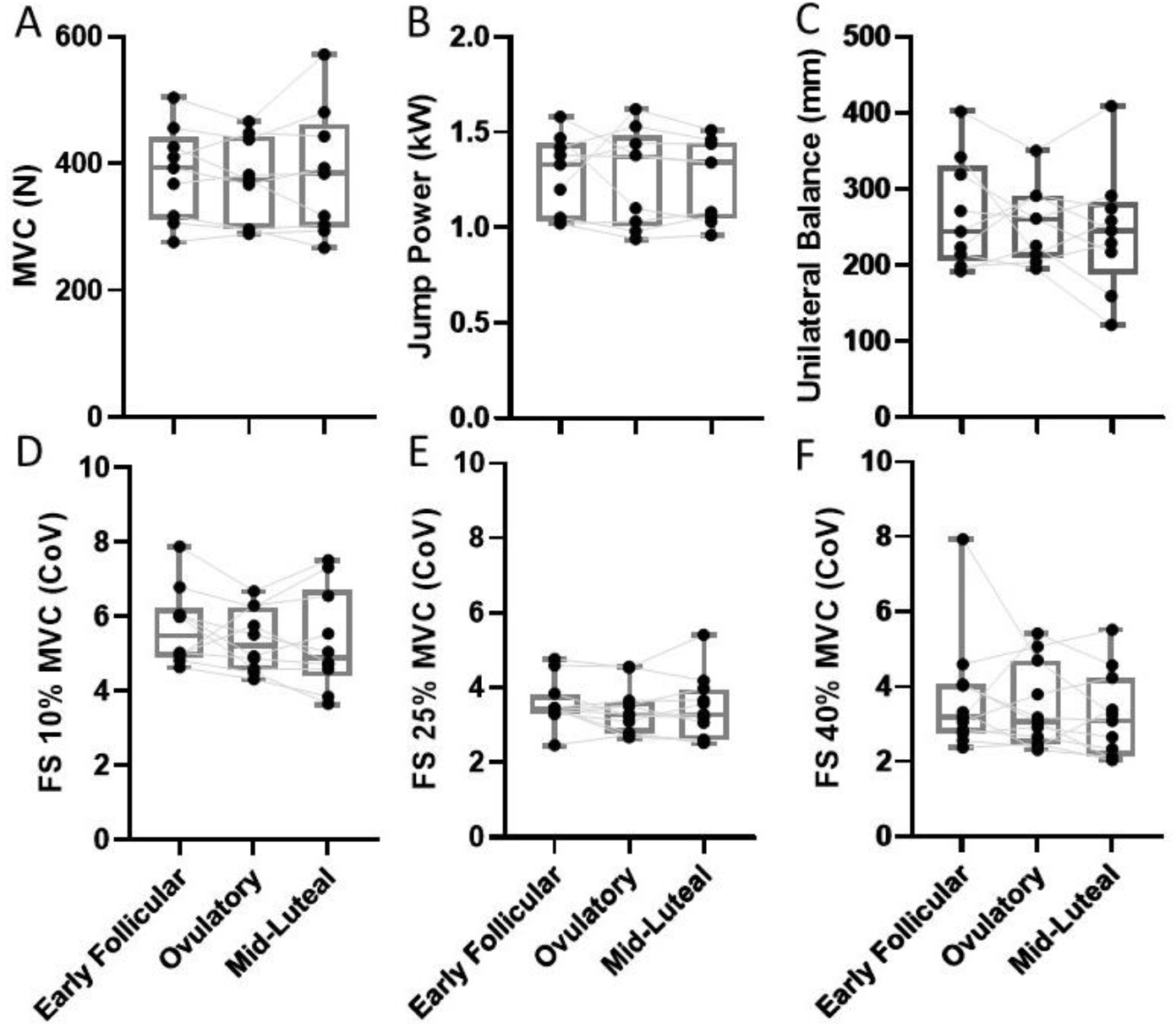
Knee extensor maximal isometric strength (A), jump power (B), right leg single balance (C), knee extensor force steadiness at 10% (A), 25% (E), 40% MVC (F), assessed at early follicular, ovulatory and mid-luteal phases of the menstrual cycle. MVC: maximum voluntary contraction, N; newtons, W; watts, mm; millimetres, FS: force steadiness, CoV; coefficient of variation.

At 10% MVC a total of 694 MUPTs were analysed in all participants, with a total of 201 (∼22 per participant) in the early follicular phase, 239 (∼26 per participant) in the ovulatory, and 254 (∼27 per participant) in the mid-luteal phase. At 25% MVC a total of 939 MUPTs were analysed in all participants, with a total of 326 (∼35 per participant) in the early follicular phase, 307 (∼36 per participant) in the ovulatory, and 306 (∼34 per participant) in the mid-luteal phase. Individual means and all MU data reflecting firing rate and firing rate variability at 10 and 25% MVC are shown in Figure 3A and B, with statistical outputs from multi-level models shown in Table 1. At 10% MVC, multi-level models revealed a lower MU firing rate at ovulation and mid luteal phases when compared to the early follicular phase (both p<0.001), and no difference between ovulation and mid luteal phases (p=0.792). Firing rate variability did not differ across any of the 3 phases. At 25% MVC, MU firing rate nor firing rate variability differed across the menstrual cycle (all p>0.3) (Table 1).

**Table 1.** Multilevel model analysis summary for motor unit parameters across three points of the menstrual cycle, at 10% and 25% MVC.

|  | 10% MVC |  |  | 25% MVC |  |  |
| --- | --- | --- | --- | --- | --- | --- |
|  | Beta | 95 CI | P value | Beta | 95 CI | P value |
| **Firing rate (Hz)** |  |  |  |  |  |  |
| *EF v Ov* | **-0.804** | **-1.206 - -0.402** | **<0.001** | 0.005 | -0.341 – 0.352 | 0.976 |
| *EF v ML* | **-0.855** | **-1.255 - -0.456** | **<0.001** | -0.110 | -0.464 – 0.245 | 0.546 |
| *Ov v ML* | -0.051 | -0.432 – 0.330 | 0.792 | -0.114 | -0.474 – 0.244 | 0.532 |
| **Firing rate Variability** |  |  |  |  |  |  |
| *EF v Ov* | 0.005 | -0.002 – 0.010 | 0.146 | -0.001 | -0.008 – 0.005 | 0.692 |
| *EF v ML* | -0.003 | -0.009 – 0.004 | 0.376 | 0.002 | -0.005 – 0.009 | 0.540 |
| *Ov v ML* | -0.007 | -0.013 – -0.001 | 0.078 | 0.003 | -0.003 – 0.010 | 0.323 |
| **MUP Area (µV.ms)** |  |  |  |  |  |  |
| *EF v Ov* | -70.79 | -142 – 0.53 | 0.059 | -27.64 | -100.8 – 45.6 | 0.460 |
| *EF v ML* | 32.19 | -39.62 – 104.1 | 0.381 | 70.44 | -3.05 – 143.9 | 0.061 |
| *Ov v ML* | **102.9** | **35.19 – 170.78** | **0.003** | **98.09** | **23.1 – 173.1** | **0.031** |
| **MUP Turns** |  |  |  |  |  |  |
| *EF v Ov* | 0.095 | -0.243 – 0.433 | 0.582 | -0.031 | -0.255 – 0.193 | 0.788 |
| *EF v ML* | **0.604** | **0.264 – 0.944** | **0.001** | **0.385** | **0.159 – 0.610** | **0.001** |
| *Ov v ML* | **0.509** | **0.188 – 0.830** | **0.003** | **0.415** | **0.186 – 0.645** | **0.001** |
| **NMJ transmission instability (%)** |  |  |  |  |  |  |
| *EF v Ov* | 0.283 | -1.94 – 2.51 | 0.804 | 0.221 | -2.01 – 2.46 | 0.846 |
| *EF v ML* | -1.029 | -3.15 – 1.09 | 0.345 | -0.475 | -2.85 – 1.35 | 0.490 |
| *Ov v ML* | -1.311 | -3.45 – 0.83 | 0.688 | -0.966 | -3.15 – 1.22 | 0.999 |

Model outputs show unstandardised beta, 95% confidence interval and associated p-values. MVC; maximum voluntary contraction, EF; early follicular, Ov; ovulation; ML; mid luteal. MUP; motor unit potential, NMJ; neuromuscular junction.

**Figure 3.**
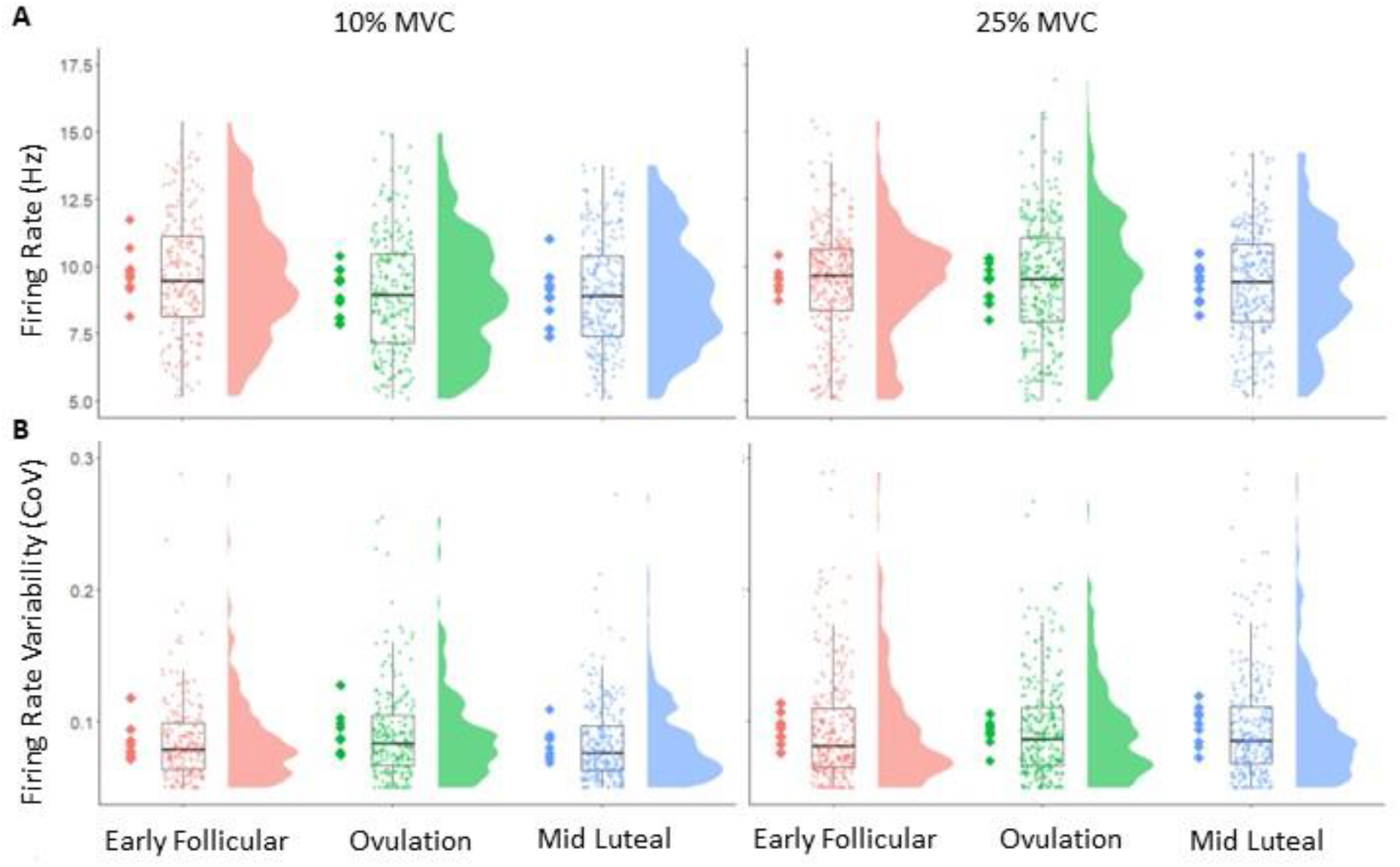
Motor unit firing rate (A) and firing rate variability (B) at 3 stages of the menstrual cycle. Data show individual participant means as diamonds, with all analysed MUPTs shown in boxplots and distribution plots. MVC; maximum voluntary contraction, CoV; coefficient of variation.

MUP area measured at 10% MVC did not differ significantly from early follicular to ovulation (p=0.059), or mid luteal (p=0.381), but was larger at mid-luteal compared to ovulation phases (p=0.003). A similar pattern was observed at 25% MVC, with no difference from early follicular to ovulation and mid-luteal (both p>0.06), but a larger MUP area at mid luteal compared to ovulation (p=0.031) (Table 1). MUP complexity, assessed via the number of turns, did not differ from early follicular to ovulation, but was more complex at mid luteal when compared to ovulation (p=0.001) and early follicular (p=0.003) phases. Again, this same pattern was observed at 25% MVC, with no change from early follicular to ovulation (p=0.788), and more complex at mid luteal when compared to ovulation (p=0.001) and early follicular (p=0.001) phases. NMJ transmission instability, measured via NFM jiggle, did not differ at any time point at 10 or 25% MVC (all p>0.3). Individual means and all MUPT data reflecting MUP area, complexity and NMJ transmission instability at 10 and 25% MVC are shown in Figure 4, with model outputs shown in Table 1. Standardised β coefficients of all MUP characteristics recorded at 10% and 25% MVC compared across timepoints are shown in Figure 5.

**Figure 4.**
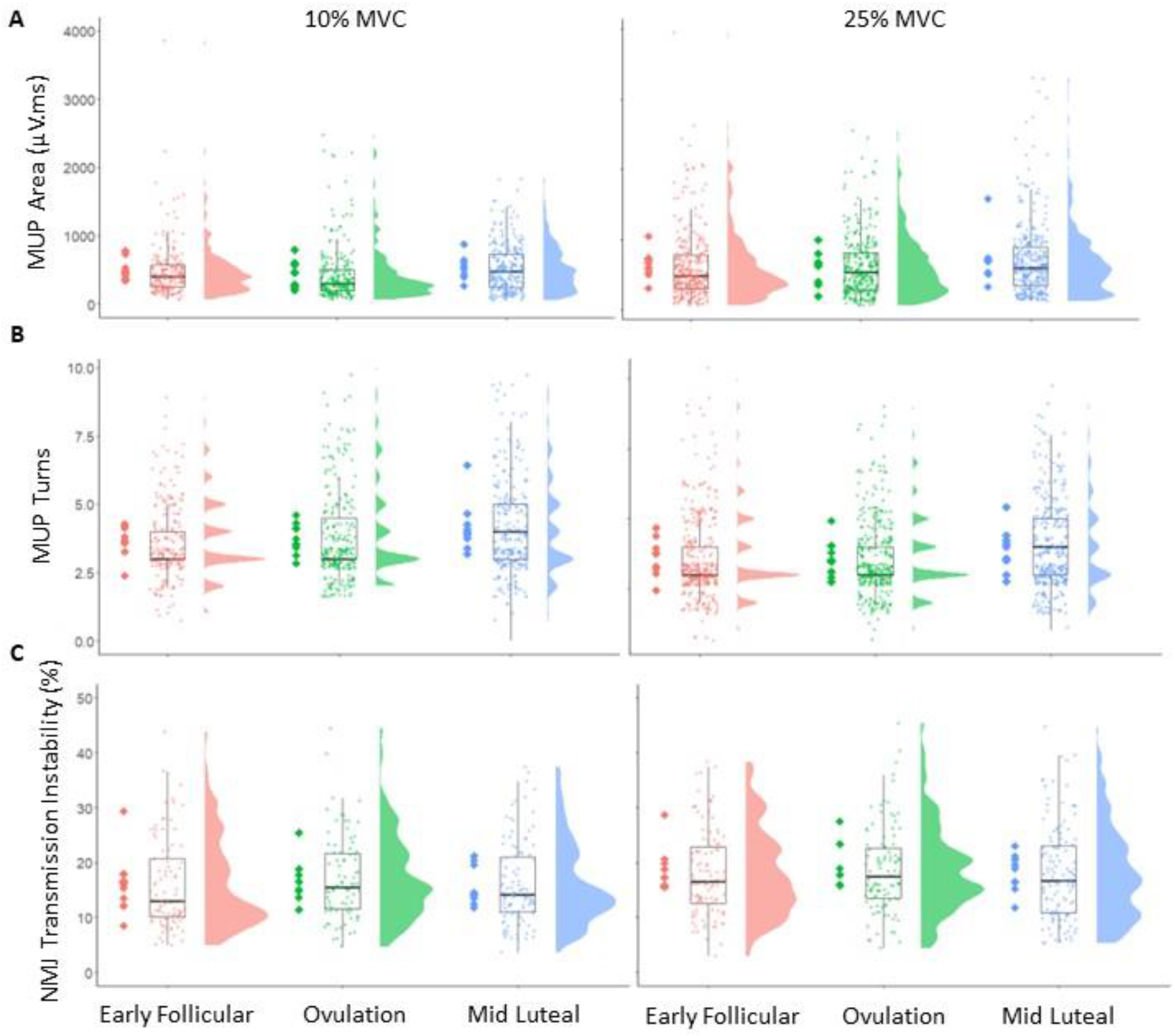
Motor unit potential area (A), complexity (number of turns) (B), and NMJ transmission instability (C) at 3 stages of the menstrual cycle. Data show individual participant means as diamonds, with all recorded MUs shown in boxplots and distribution plots. MUP; motor unit potential, µV.ms; microvolt millisecond, NMJ; neuromuscular junction.

**Figure 5.**
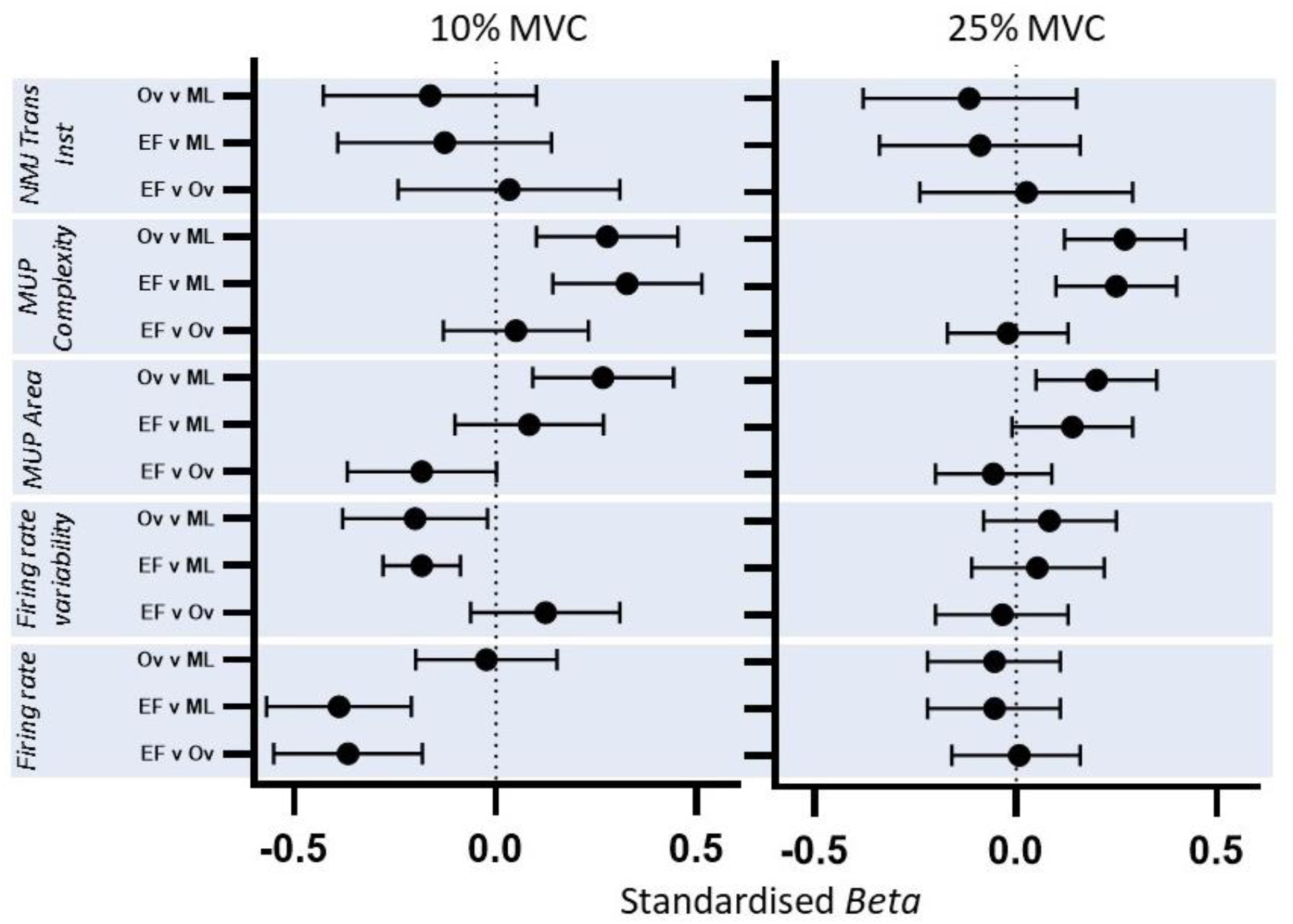
Standardised beta coefficients comparing motor unit parameters recorded during 10% and 25% MVC contractions. Beta value and 95% confidence intervals (CI) represent the standardized model predicted change per unit between menstrual cycle phases. MUP; motor unit potential, NMJ; neuromuscular junction. EF; early follicular, Ov; ovulation, ML; mid luteal.

We next explored recruitment strategies when moving between contraction levels (10% to 25% MVC) at each timepoint in the cycle. There were no statistically significant differences in MU firing rate at early follicular phase (p=0.179), however firing rate increased from 10 to 25% MVC at ovulation (p=0.047) and mid-luteal phase (p=0.015). There was a significant contraction level x time interaction when comparing the early follicular phase with the ovulation (p=0.09) and the mid luteal phase (p=0.009), indicating a larger contraction level related increase at these time points (Table 2). Firing rate variability increased in the early follicular (p=0.004) and mid luteal phase (p<0.001), with no change in the ovulation phase (p=0.290) (Table 2). There were no statistically significant contraction level x time interactions in firing rate variability. MUP area increased with increased contraction level at all time points across the cycle (all p<0.002) with no interactions, indicating the degree to which MUP area increased from 10% to 25% MVC did not differ at each timepoint. The number of MUP turns did not differ between 10% and 25% MVC contractions at any time point (all p>0.07). NMJ transmission instability increased with larger contractions at all timepoints (all p<0.049), with no contraction level x time interactions (Table 2). Standardised β coefficients of all MUP characteristics displaying relative differences between 10% and 25% MVC at each timepoint are shown in Figure 6.

**Table 2.** Multilevel model analysis summary comparing motor unit parameters from 10% to 25% MVC, at 3 timepoints of the menstrual cycle.

|  | Beta | 95 CI | P value |
| --- | --- | --- | --- |
| **Firing rate (Hz)** |  |  |  |
| *Early follicular* | -0.266 | -0.654 – 0.121 | 0.179 |
| *Ovulation* | **0.380** | **-0.010 – 0.770** | **0.047** |
| *Mid luteal* | **0.434** | **0.085 – 0.784** | **0.015** |
| **Firing rate Variability** |  |  |  |
| *Early follicular* | **0.010** | **0.003 – 0.017** | **0.004** |
| *Ovulation* | 0.004 | -0.003 – 0.010 | 0.290 |
| *Mid luteal* | **0.013** | **0.007 – 0.019** | **<0.001** |
| **MUP Area (μV.ms)** |  |  |  |
| *Early follicular* | **135.21** | **56.47 – 213.95** | **0.001** |
| *Ovulation* | **202.84** | **136.48 – 269.21** | **<0.001** |
| *Mid luteal* | **202.65** | **131.92 – 273.38** | **<0.001** |
| **MUP Turns** |  |  |  |
| *Early follicular* | -0.003 | -0.253 – 0.246 | 0.980 |
| *Ovulation* | -0.093 | -0.346 – -0.716 | 0.474 |
| *Mid luteal* | -0.266 | -0.560 – -1.769 | 0.077 |
| **NMJ transmission instability (%)** |  |  |  |
| *Early follicular* | **2.114** | **0.064 – 4.164** | **0.045** |
| *Ovulation* | **2.005** | **-0.087 – 4.096** | **0.049** |
| *Mid luteal* | **2.221** | **0.096 – 4.345** | **0.043** |

Model outputs show unstandardised beta, 95% confidence interval and associated p-values. MVC; maximum voluntary contraction. MUP; motor unit potential, NMJ; neuromuscular junction.

**Figure 6.**
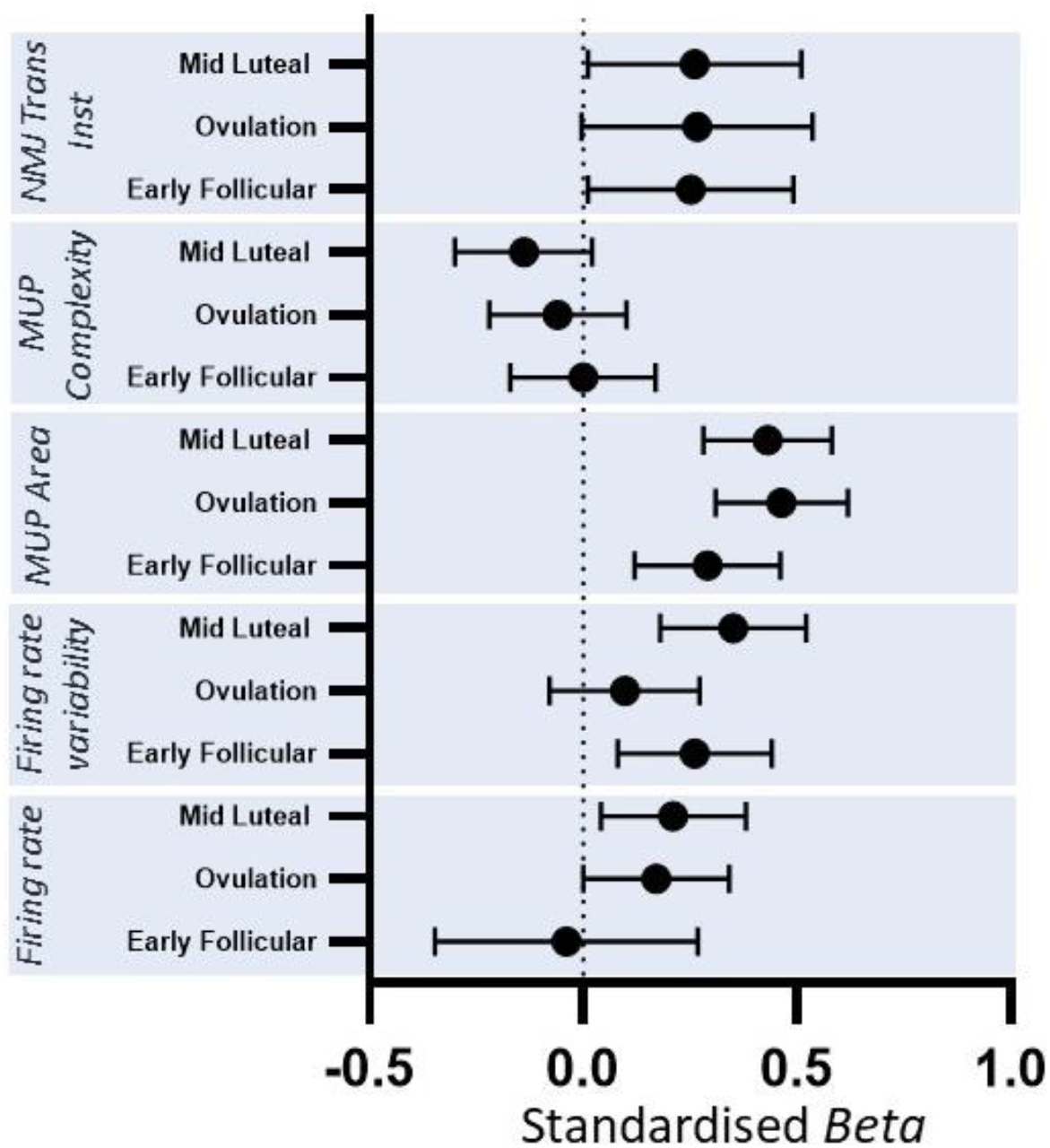
Standardised beta coefficients displaying MUPT parameter differences from 10% to 25% MVC at each timepoint of the cycle. Beta value and 95% confidence intervals (CI) represent the standardized model predicted change per unit between 10% and 25% MVC at each phase of the menstrual cycle. MUP; motor unit potential, NMJ; neuromuscular junction. EF; early follicular, Ov; ovulation, ML; mid luteal.

We then explored the relationship between each hormone and each mean MUPT parameter across timepoints. Regression models with time as a covariate revealed no significant association between circulating concentrations of estrogen and MUPT parameters measured at 10% or 25% MVC (all p>0.1). There were also no significant associations with progesterone and MUPT parameters measured at 10% and 25% MVC (p>0.17).

## Discussion

This study sought to determine the potential alteration of neuromuscular performance and individual VL MU function across the menstrual cycle in eumenorrheic females and to assess the influence of hormonal fluctuations on these measures. Most notably, low threshold MU firing rate was lower in the latter two phases of the cycle, matched by an increase in MUP size in the final phase which coincided with an increase in circulating progesterone. Multiple measures of performance relative to neuromuscular strength and control did not differ at any time point of the cycle. Although statistical outputs revealed no clear relationship between hormone concentrations and MU function, the suppression of MU firing rate in the presence of increased progesterone suggests a neuroinhibitory effect across the latter half of the menstrual cycle preferentially affecting early recruited MUs.

Neuromuscular performance was measured through maximum voluntary contraction, jump power, single leg balance, and force steadiness. Our data did not identify any differences across these measures which aligns with recent reports of very limited performance decrements across the menstrual cycle of eumenorrheic women (McNulty *et al*., 2020). Yet it is still quite frequently noted that females report declines in neuromuscular performance through a myriad of symptoms at varying times of the menstrual cycle (Chantler, Mitchell and Fuller, 2009; Findlay *et al*., 2020). Although the functionality data presented herein cannot support this, the range of physiological performance assessments is vast, and decrements may be noted in less controlled or more prolonged assessments to those applied here. Moreover, the large heterogeneity of the menstrual cycle and its effects, reflected in the known inter-individual and intra-individual variations in outcomes of the menstrual cycle e.g. length, bleeding patterns, symptoms and severity (Bull *et al*., 2019; Bruinvels *et al*., 2021b), and a possible disconnect between circulating hormones and their respective functional receptors may differentially influence outcomes of performance assessments.

MU firing rate at 10% MVC was highest at the early follicular phase and decreased in ovulation and mid luteal phases, with no change across the cycle at 25% MVC indicating these effects are clearly contraction level specific. This difference at 10% MVC is comparatively large; it is greater than that we reported in the VL of young males following 15 days of limb immobilisation at normalised and absolute contraction intensities (Inns *et al*., 2022), and is larger than that reported following 4 week resistance training in the tibialis anterior (Del Vecchio *et al*., 2019). However, in these studies the neuromodulatory alterations were aligned with changes in force production as a result of the intervention, which was not identified in this female cohort which lacked an intervention. These current findings are also in slight contrast to those of Tenan et al (2013), who reported an increase in firing rate of vastus medialis MUs across five phases of the menstrual cycle quantified via basal body temperature mapping. When comparing the early follicular to the mid-luteal phases, the mean initial firing rate increased from 8.4 to 9.1 Hz in 140 individual MUs recorded from 7 females (Tenan *et al*., 2013). Although a different muscle to that assessed here, VL and VM have similar firing rates (Avrillon *et al*., 2021) and both share common synaptic input (Rossato *et al*., 2022). It is not entirely clear why the two synergist muscles may be differentially affected by menstrual cycle stage, although some reasoning could be inferred via the number of MUPs within a MUPT used to calculate MU firing rate; all observed within a MUPT of the sustained plateau of a 12-15s contraction in the current study, compared to the average of the first 3 of a ramped contraction (Tenan *et al*., 2013).

Firing rate is a key determinant of muscle force and typically increases with increased contraction intensity (Orssatto *et al*., 2021; Guo *et al*., 2022). The difference between 10 and 25% contractions here was more pronounced in the latter phases, again pointing to a suppression of firing rate of early recruited MUs in the presence of progesterone, which as outlined above, is largely inhibitory. However, these findings also negate the potential net excitatory effects of estrogen (Smith and Woolley, 2004; Mukherjee *et al*., 2017; Del Río *et al*., 2018) and suggest progesterone is the more dominant of the two in this respect. MU firing rate is mediated by ionotropic input from descending pathways, peripheral afferents and spinal interneurons, and via neuromodulatory inputs that can alter the intrinsic excitability of the motoneuron through voltage-dependent ion channels. These latter inputs are known as persistent inward currents (PICs) and act to amplify and prolong synaptic inputs in a non-linear fashion (Heckman, Gorassini and Bennett, 2005). Lower threshold MUs are highly sensitive to inhibitory input (Lee and Heckman, 1998) and it is possible they are more dependent on prolonged PIC activity, which increases with increased contraction level (Orssatto *et al*., 2021), than on descending inputs. As such, they may be more susceptible to the inhibitory effects of progesterone. A variable response of MUs across the entire pool is not uncommon as lower threshold MUs are also more susceptible to pain induced inhibition (Martinez-Valdes *et al*., 2020), and although differing to the contraction levels applied here, this likely results from a non-uniform distribution of inhibitory inputs across the MU pool (Mesquita, Škarabot and Pearcey, 2020). Firing rate variability partly influences force steadiness (Moritz *et al*., 2005; Ely *et al*., 2022) and this did not differ across the cycle at either contraction level, aligned with no difference of force steadiness measures.

The area of a MUP typically increases with larger contraction levels as a result of the recruitment of larger MUs (Guo *et al*., 2022). A similar pattern was observed here as increases in MUP area from low to mid-contractions occurred to a similar extent at each phase of the cycle. When comparing across the cycle, MUP area was greatest in the mid luteal phase at both contraction levels. We have previously reported that at a given contraction level, females have smaller MUs than males but possibly compensate for this with higher comparative firing rates (Guo *et al*., 2022). The same may be true here, where MUP area is greater when firing rate is lowest, indicating a minor alteration of recruitment strategy during the mid-luteal phase, compensating for reductions in firing rate at the same relative contraction intensity. However, the difference in MUP area was not observed at the ovulation phase where firing rate was also comparatively lower. We observed a greater number of MUP turns, or MUP complexity in the mid luteal phase where MUP area was also the largest. When the propagation of action potentials along muscle fibres of the same MU are temporally dispersed, the complexity, or number of turns of the recorded MUP, is greater (Piasecki, Garnés-Camarena and Stashuk, 2021). This can increase acutely in humans following disuse periods of 10 (Sarto *et al*., 2022) and 15 days (Inns *et al*., 2022), occurring in response to pre- and/or post-synaptic mechanisms; a result of temporally dispersed yet consistent transmission at the NMJ of MU fibres, or propagation along MU fibres. Estrogen-induced inhibition of voltage-gated Ca^2+^ channels occurs in smooth muscle (Wray and Noble, 2008), and although the current data are far from definitive in this aspect, similar effects at the NMJ could result in increased MUP complexity as transmission at fibres of the same MU becomes more asynchronous. NMJ transmission instability estimated by iEMG and near fibre analysis has consistently shown an increased instability of aged human NMJ transmission (Hourigan *et al*., 2015; Piasecki *et al*., 2016, 2020) and feasibly contributes to age-related decreases in strength. The lack of a difference across the menstrual cycle shown here does not support acute alterations at the NMJ that may be influenced by hormonal fluctuations or other cycle-associated adaptation.

### Strengths and limitations

To our knowledge, this is the first study to report multiple aspects of neuromuscular and individual MU adaptations of the vastus lateralis muscle across three timepoints of the menstrual cycle with quantification of plasma estrogen and progesterone. The iEMG techniques applied enable identification of adaptations occurring centrally (MU firing rate and firing rate variability) alongside those occurring peripherally at the NMJ and muscle fibre. We applied rigorous methods to accurately quantify the cycle period via self-assessment and hormone quantification and obtained all data within a single cycle to avoid multi-cycle variability. This latter point may be viewed by some as a limitation; all participants completed the three assessments in phase order from the early follicular, to ovulation and finally to the mid-luteal phase which presents the possibility of a learning effect. However, we anticipate this to be minimal given the lack of difference in any performance measure and there were at least seven days between our chosen phases of the cycle, further minimising any possible learning effect.

### Conclusion

Knee extensor muscle force, power, and force steadiness is unaltered across the female menstrual cycle under conditions of fluctuating estrogen and progesterone. There is evidence of adaptation of MU firing rate of low threshold MUs and a probable alteration of recruitment strategy to achieve a given force. This occurred alongside higher levels of progesterone which may suggest an inhibitory effect of lower threshold MUs, whilst NMJ transmission instability was unaltered and highlights minimal effects of the cycle on the peripheral motor system. However, although notably interesting from a physiological perspective, these observations had no clear influence on any of the functional assessments of neuromuscular strength and control. Furthermore, the fluctuations in estrogen and progesterone could not statistically explain any of the observed MU adaptations and may in fact be a result of the combined myriad of physiological adaptations that occur across the cycle independent of these hormones. Collectively, our human data reveal minimal effects of the menstrual cycle on neuromuscular performance with a probable suppression of firing rate early recruited motor units with increased progesterone. These findings can inform upon the effective control of stages of the menstrual cycle, where required, and minimize the perceived limitation of including females in neuromuscular physiology studies, ultimately supporting their greater inclusion.

## Data Availability Statement

The datasets generated and analysed during the current study are available from the corresponding author upon reasonable request.

## Acknowledgements

We are grateful to Dr Paul Ansdell for his critical insight to the physiology of the menstrual cycle.

## References

Ansdell, P. et al. (2019) ‘Menstrual cycle-associated modulations in neuromuscular function and fatigability of the knee extensors in eumenorrheic women’, Journal of Applied Physiology, 126(6). Available at: https://doi.org/10.1152/japplphysiol.01041.2018.

Avrillon, S. et al. (2021) ‘Individual differences in the neural strategies to control the lateral and medial head of the quadriceps during a mechanically constrained task’, Journal of Applied Physiology, 130(1), pp. 269–281. Available at: https://doi.org/10.1152/japplphysiol.00653.2020.

Barth, C., Villringer, A. and Sacher, J. (2015) ‘Sex hormones affect neurotransmitters and shape the adult female brain during hormonal transition periods’, Frontiers in Neuroscience, 9, p. 37. Available at: https://doi.org/10.3389/fnins.2015.00037.

Bates, D. et al. (2015) ‘Fitting Linear Mixed-Effects Models Using lme4’, Journal of Statistical Software, 67(1). Available at: https://doi.org/10.18637/jss.v067.i01.

Bethea, C.L. and Reddy, A.P. (2012) ‘Ovarian steroids increase glutamatergic related gene expression in serotonin neurons of macaques’, Molecular and Cellular Neurosciences, 49(3), pp. 251–262. Available at: https://doi.org/10.1016/j.mcn.2011.11.005.

Birch, K. and Reilly, T. (2002) ‘The diurnal rhythm in isometric muscular performance differs with eumenorrheic menstrual cycle phase’, Chronobiology International, 19(4). Available at: https://doi.org/10.1081/CBI-120006083.

Bitran, D., Shiekh, M. and McLeod, M. (1995) ‘Anxiolytic effect of progesterone is mediated by the neurosteroid allopregnanolone at brain GABAA receptors’, Journal of Neuroendocrinology, 7(3), pp. 171–177. Available at: https://doi.org/10.1111/j.1365-2826.1995.tb00744.x.

Blagrove, R.C., Bruinvels, G. and Pedlar, C.R. (2020) ‘Variations in strength-related measures during the menstrual cycle in eumenorrheic women: A systematic review and meta-analysis’, Journal of Science and Medicine in Sport, 23(12). Available at: https://doi.org/10.1016/j.jsams.2020.04.022.

Bruinvels, G. et al. (2016) ‘The prevalence and impact of heavy menstrual bleeding (Menorrhagia) in elite and non-elite athletes’, PLoS ONE, 11(2). Available at: https://doi.org/10.1371/journal.pone.0149881.

Bruinvels, G. et al. (2021a) ‘Prevalence and frequency of menstrual cycle symptoms are associated with availability to train and compete: A study of 6812 exercising women recruited using the Strava exercise app’, British Journal of Sports Medicine, 55(8). Available at: https://doi.org/10.1136/bjsports-2020-102792.

Bruinvels, G. et al. (2021b) ‘Prevalence and frequency of menstrual cycle symptoms are associated with availability to train and compete: a study of 6812 exercising women recruited using the Strava exercise app’, British Journal of Sports Medicine, 55(8), pp. 438–443. Available at: https://doi.org/10.1136/bjsports-2020-102792.

Bruinvels, G. et al. (2022) ‘How Lifestyle Changes during the COVID-19 Global Pandemic Affected the Pattern and Symptoms of the Menstrual Cycle’, International Journal of Environmental Research and Public Health, 19(20), p. 13622. Available at: https://doi.org/10.3390/ijerph192013622.

Bull, J.R. et al. (2019) ‘Real-world menstrual cycle characteristics of more than 600,000 menstrual cycles’, NPJ digital medicine, 2, p. 83. Available at: https://doi.org/10.1038/s41746-019-0152-7.

Chantler, I., Mitchell, D. and Fuller, A. (2009) ‘Diclofenac Potassium Attenuates Dysmenorrhea and Restores Exercise Performance in Women With Primary Dysmenorrhea’, The Journal of Pain, 10(2), pp. 191–200. Available at: https://doi.org/10.1016/j.jpain.2008.08.006.

Davies, B.N., Elford, J.C. and Jamieson, K.F. (1991) ‘Variations in performance in simple muscle tests at different phases of the menstrual cycle’, The Journal of Sports Medicine and Physical Fitness, 31(4), pp. 532–537.

Del Río, J.P. et al. (2018) ‘Steroid Hormones and Their Action in Women’s Brains: The Importance of Hormonal Balance’, Frontiers in Public Health, 6. Available at: https://www.frontiersin.org/articles/10.3389/fpubh.2018.00141 x(Accessed: 17 January 2023).

Del Vecchio, A. et al. (2019) ‘The increase in muscle force after 4 weeks of strength training is mediated by adaptations in motor unit recruitment and rate coding’, The Journal of Physiology, 597(7), pp. 1873–1887. Available at: https://doi.org/10.1113/JP277250.

Dibrezzo, R., Fort, I.L. and Brown, B. (1988) ‘Dynamic strength and work variations during three stages of the menstrual cycle’, Journal of Orthopaedic and Sports Physical Therapy, 10(4). Available at: https://doi.org/10.2519/jospt.1988.10.4.113.

Ekenros, L. et al. (2017) ‘Expression of sex steroid hormone receptors in human skeletal muscle during the menstrual cycle’, Acta Physiologica, 219(2), pp. 486–493. Available at: https://doi.org/10.1111/apha.12757.

Elliott, K.J. et al. (2003) ‘Effect of menstrual cycle phase on the concentration of bioavailable 17-β oestradiol and testosterone and muscle strength’, Clinical Science, 105(6). Available at: https://doi.org/10.1042/CS20020360.

Ely, I.A. et al. (2022) ‘Training-induced improvements in knee extensor force accuracy are associated with reduced vastus lateralis motor unit firing variability’, Experimental Physiology, 107(9), pp. 1061–1070. Available at: https://doi.org/10.1113/EP090367.

Eshima, J. et al. (2019) ‘Monitoring changes in the healthy female metabolome across the menstrual cycle using GC × GC-TOFMS’, Journal of Chromatography B, 1121, pp. 48–57. Available at: https://doi.org/10.1016/j.jchromb.2019.04.046.

Fehring, R.J., Schneider, M. and Raviele, K. (2006) ‘Variability in the Phases of the Menstrual Cycle’, Journal of Obstetric, Gynecologic & Neonatal Nursing, 35(3), pp. 376–384. Available at: https://doi.org/10.1111/j.1552-6909.2006.00051.x.

Findlay, R.J. et al. (2020) ‘How the menstrual cycle and menstruation affect sporting performance: experiences and perceptions of elite female rugby players’, British Journal of Sports Medicine, 54(18), pp. 1108–1113. Available at: https://doi.org/10.1136/bjsports-2019-101486.

Giovannini, F. et al. (2002) ‘Calcium channel subtypes contributing to acetylcholine release from normal, 4-aminopyridine-treated and myasthenic syndrome auto-antibodies-affected neuromuscular junctions: Calcium channels at the mouse neuromuscular junction’, British Journal of Pharmacology, 136(8), pp. 1135–1145. Available at: https://doi.org/10.1038/sj.bjp.0704818.

Greeves, J.P. et al. (1997) ‘Effects of acute changes in oestrogen on muscle function of the first dorsal interosseus muscle in humans’, The Journal of Physiology, 500 (Pt 1)(Pt 1), pp. 265–270. Available at: https://doi.org/10.1113/jphysiol.1997.sp022016.

Guo, Y. et al. (2022) ‘Neuromuscular recruitment strategies of the vastus lateralis according to sex’, Acta Physiologica [Preprint]. Available at: https://doi.org/10.1111/apha.13803.

Heckman, C.J., Gorassini, M.A. and Bennett, D.J. (2005) ‘Persistent inward currents in motoneuron dendrites: Implications for motor output’, Muscle & Nerve, 31(2), pp. 135–156. Available at: https://doi.org/10.1002/mus.20261.

Hourigan, M.L. et al. (2015) ‘Increased motor unit potential shape variability across consecutive motor unit discharges in the tibialis anterior and vastus medialis muscles of healthy older subjects’, Clinical Neurophysiology, 126(12), pp. 2381–2389. Available at: https://doi.org/10.1016/j.clinph.2015.02.002.

Hsu, F.C. and Smith, S.S. (2003) ‘Progesterone withdrawal reduces paired-pulse inhibition in rat hippocampus: Dependence on GABAAreceptor α4 subunit upregulation’, Journal of Neurophysiology, 89(1). Available at: https://doi.org/10.1152/jn.00195.2002.

Inns, T.B. et al. (2022) ‘Motor unit dysregulation following 15 days of unilateral lower limb immobilisation’, The Journal of Physiology, 600(21), pp. 4753–4769. Available at: https://doi.org/10.1113/JP283425.

Janse De Jonge, X.A.K. et al. (2001) ‘The influence of menstrual cycle phase on skeletal muscle contractile characteristics in humans’, Journal of Physiology, 530(1). Available at: https://doi.org/10.1111/j.1469-7793.2001.0161m.x.

Jones, E.J. et al. (2021) ‘Lifelong exercise is associated with more homogeneous motor unit potential features across deep and superficial areas of vastus lateralis’, GeroScience, 43(4). Available at: https://doi.org/10.1007/s11357-021-00356-8.

de Jonge, X.J., Thompson, B. and Ahreum, H.A.N. (2019) ‘Methodological Recommendations for Menstrual Cycle Research in Sports and Exercise’, Medicine and Science in Sports and Exercise, 51(12). Available at: https://doi.org/10.1249/MSS.0000000000002073.

Kubo, K. et al. (2009) ‘Muscle and tendon properties during menstrual cycle’, International Journal of Sports Medicine, 30(2). Available at: https://doi.org/10.1055/s-0028-1104573.

Lee, R.H. and Heckman, C.J. (1998) ‘Bistability in Spinal Motoneurons In Vivo: Systematic Variations in Persistent Inward Currents’, Journal of Neurophysiology, 80(2), pp. 583–593. Available at: https://doi.org/10.1152/jn.1998.80.2.583.

Lowe, D.A., Baltgalvis, K.A. and Greising, S.M. (2010) ‘Mechanisms Behind Estrogen’s Beneficial Effect on Muscle Strength in Females’, Exercise and Sport Sciences Reviews, 38(2), pp. 61–67. Available at: https://doi.org/10.1097/JES.0b013e3181d496bc.

Martin, D. et al. (2018) ‘Period prevalence and perceived side effects of hormonal contraceptive use and the menstrual cycle in elite athletes’, International Journal of Sports Physiology and Performance, 13(7). Available at: https://doi.org/10.1123/ijspp.2017-0330.

Martin, D. et al. (2021) ‘Injury Incidence Across the Menstrual Cycle in International Footballers’, Frontiers in Sports and Active Living, 3. Available at: https://www.frontiersin.org/articles/10.3389/fspor.2021.616999 x(Accessed: 17 January 2023).

Martinez-Valdes, E. et al. (2020) ‘Divergent response of low-versus high-threshold motor units to experimental muscle pain’, The Journal of Physiology, 598(11), pp. 2093–2108. Available at: https://doi.org/10.1113/JP279225.

McNulty, K.L. et al. (2020) ‘The Effects of Menstrual Cycle Phase on Exercise Performance in Eumenorrheic Women: A Systematic Review and Meta-Analysis’, Sports Medicine, 50(10). Available at: https://doi.org/10.1007/s40279-020-01319-3.

Melcangi, R.C. et al. (2014) ‘Levels and actions of progesterone and its metabolites in the nervous system during physiological and pathological conditions’, Progress in Neurobiology, 113, pp. 56–69. Available at: https://doi.org/10.1016/j.pneurobio.2013.07.006.

Mesquita, R.N.O., Škarabot, J. and Pearcey, G.E.P. (2020) ‘Low-threshold motor units can be a pain during experimental muscle pain’, The Journal of Physiology, 598(13), pp. 2545–2547. Available at: https://doi.org/10.1113/JP279872.

Messinis, I.E., Messini, C.I. and Dafopoulos, K. (2014) ‘Novel aspects of the endocrinology of the menstrual cycle’, Reproductive BioMedicine Online, 28(6). Available at: https://doi.org/10.1016/j.rbmo.2014.02.003.

Michopoulos, V., Berga, S.L. and Wilson, M.E. (2011) ‘Estradiol and progesterone modify the effects of the serotonin reuptake transporter polymorphism on serotonergic responsivity to citalopram’, Experimental and Clinical Psychopharmacology, 19(6), pp. 401–408. Available at: https://doi.org/10.1037/a0025008.

Moritz, C.T. et al. (2005) ‘Discharge rate variability influences the variation in force fluctuations across the working range of a hand muscle’, Journal of Neurophysiology, 93(5), pp. 2449–2459. Available at: https://doi.org/10.1152/jn.01122.2004.

Mukherjee, J. et al. (2017) ‘Estradiol modulates the efficacy of synaptic inhibition by decreasing the dwell time of GABAA receptors at inhibitory synapses’, Proceedings of the National Academy of Sciences, 114(44), pp. 11763–11768. Available at: https://doi.org/10.1073/pnas.1705075114.

Orssatto, L.B.R. et al. (2021) ‘Estimates of persistent inward currents increase with the level of voluntary drive in low-threshold motor units of plantar flexor muscles’, Journal of Neurophysiology, 125(5), pp. 1746–1754. Available at: https://doi.org/10.1152/jn.00697.2020.

Phillips, S.K. et al. (1996) ‘Changes in maximal voluntary force of human adductor pollicis muscle during the menstrual cycle’, The Journal of Physiology, 496 (Pt 2)(Pt 2), pp. 551–557. Available at: https://doi.org/10.1113/jphysiol.1996.sp021706.

Piasecki, J. et al. (2020) ‘Influence of sex on the age-related adaptations of neuromuscular function and motor unit properties in elite masters athletes’, Journal of Physiology [Preprint]. Available at: https://doi.org/10.1113/JP280679.

Piasecki, M. et al. (2016) ‘Age-related neuromuscular changes affecting human vastus lateralis’, Journal of Physiology, 594(16). Available at: https://doi.org/10.1113/JP271087.

Piasecki, M., Garnés-Camarena, O. and Stashuk, D.W. (2021) ‘Near-fiber electromyography’, Clinical Neurophysiology, 132(5), pp. 1089–1104. Available at: https://doi.org/10.1016/j.clinph.2021.02.008.

Potier, M. et al. (2016) ‘Temporal Memory and Its Enhancement by Estradiol Requires Surface Dynamics of Hippocampal CA1 N-Methyl-D-Aspartate Receptors’, Biological Psychiatry, 79(9), pp. 735–745. Available at: https://doi.org/10.1016/j.biopsych.2015.07.017.

Rossato, J. et al. (2022) ‘Less common synaptic input between muscles from the same group allows for more flexible coordination strategies during a fatiguing task’, Journal of Neurophysiology, 127(2), pp. 421–433. Available at: https://doi.org/10.1152/jn.00453.2021.

Sarto, F. et al. (2022) ‘Effects of short-term unloading and active recovery on human motor unit properties, neuromuscular junction transmission and transcriptomic profile’, The Journal of Physiology, 600(21), pp. 4731–4751. Available at: https://doi.org/10.1113/JP283381.

Sarwar, R., Niclos, B.B. and Rutherford, O.M. (1996) ‘Changes in muscle strength, relaxation rate and fatiguability during the human menstrual cycle’, Journal of Physiology, 493(1). Available at: https://doi.org/10.1113/jphysiol.1996.sp021381.

Schmalenberger, K.M. et al. (2019) ‘A Systematic Review and Meta-Analysis of Within-Person Changes in Cardiac Vagal Activity across the Menstrual Cycle: Implications for Female Health and Future Studies’, Journal of Clinical Medicine, 8(11), p. 1946. Available at: https://doi.org/10.3390/jcm8111946.

Shayani, D.R. et al. (2020) ‘The role of health anxiety in the experience of perceived stress across the menstrual cycle’, Anxiety, Stress, & Coping, 33(6), pp. 706–715. Available at: https://doi.org/10.1080/10615806.2020.1802434.

Shultz, S.J. et al. (2004) ‘Relationship between sex hormones and anterior knee laxity across the menstrual cycle’, Medicine and Science in Sports and Exercise, 36(7), pp. 1165–1174. Available at: https://doi.org/10.1249/01.mss.0000132270.43579.1a.

Smith, S.S. et al. (1987) ‘Progesterone alters GABA and glutamate responsiveness: a possible mechanism for its anxiolytic action’, Brain Research, 400(2), pp. 353–359. Available at: https://doi.org/10.1016/0006-8993(87)90634-2.

Smith, S.S. (1994) ‘Female sex steroid hormones: from receptors to networks to performance--actions on the sensorimotor system’, Progress in Neurobiology, 44(1), pp. 55–86. Available at: https://doi.org/10.1016/0301-0082(94)90057-4.

Smith, S.S., Woodward, D.J. and Chapin, J.K. (1989) ‘Sex steroids modulate motor-correlated increases in cerebellar discharge’, Brain Research, 476(2). Available at: https://doi.org/10.1016/0006-8993(89)91251-1.

Smith, S.S. and Woolley, C.S. (2004) ‘Cellular and molecular effects of steroid hormones on CNS excitability’, Cleveland Clinic Journal of Medicine, 71(SUPPL. 2). Available at: https://doi.org/10.3949/ccjm.71.suppl_2.s4.

Stoffel-Wagner, B. (2001) ‘Neurosteroid metabolism in the human brain’, European Journal of Endocrinology, 145(6), pp. 669–679. Available at: https://doi.org/10.1530/eje.0.1450669.

Tenan, M.S. et al. (2013) ‘Menstrual cycle mediates vastus medialis and vastus medialis oblique muscle activity’, Medicine and Science in Sports and Exercise, 45(11). Available at: https://doi.org/10.1249/MSS.0b013e318299a69d.

Tenan, M.S., Hackney, A.C. and Griffin, L. (2016) ‘Maximal force and tremor changes across the menstrual cycle’, European Journal of Applied Physiology, 116(1). Available at: https://doi.org/10.1007/s00421-015-3258-x.

Weidauer, L. et al. (2020) ‘Neuromuscular performance changes throughout the menstrual cycle in physically active females’, Journal of Musculoskeletal & Neuronal Interactions, 20(3), pp. 314–324.

Wojtys, E.M. et al. (1998) ‘Association Between the Menstrual Cycle and Anterior Cruciate Ligament Injuries in Female Athletes’, The American Journal of Sports Medicine, 26(5), pp. 614–619. Available at: https://doi.org/10.1177/03635465980260050301.

Wray, S. and Noble, K. (2008) ‘Sex Hormones and Excitation–Contraction Coupling in the Uterus: The Effects of Oestrous and Hormones’, Journal of Neuroendocrinology, 20(4), pp. 451–461. Available at: https://doi.org/10.1111/j.1365-2826.2008.01665.x.

